# Volumetric Semantic Instance Segmentation of the Plasma Membrane of HeLa Cells

**DOI:** 10.1101/2021.04.30.442156

**Authors:** Cefa Karabağ, Martin L. Jones, Constantino Carlos Reyes-Aldasoro

## Abstract

In this work, the unsupervised volumetric semantic segmentation of the plasma membrane of HeLa cells as observed with Serial Block Face Scanning Electron Microscopy is described. The resin background of the images was segmented at different slices of a 3D stack of 518 slices with 8, 192 × 8, 192 pixels each. The background was used to create a distance map which helped identify and rank the cells by their size at each slice. The centroids of the cells detected at different slices were linked to identify them as a single cell that spanned a number of slices. A subset of these cells, i.e., largest ones and those not close to the edges were selected for further processing. The selected cells were then automatically cropped to smaller regions of interest of 2, 000 × 2, 000 × 300 voxels that were treated as cell instances. Then, for each of these volumes the nucleus was segmented and the cell was separated from any neighbouring cells through a series of traditional image processing steps that followed the plasma membrane. The segmentation process was repeated for all the regions selected. For one cell for which the ground truth was available, the algorithm provided excellent results in Accuracy (AC) and Jaccard Index (JI): Nucleus: JI = 0.9665, AC= 0.9975, Cell and Nucleus JI = 0.8711, AC = 0.9655, Cell only JI = 0.8094, AC = 0.9629. A limitation of the algorithm for the plasma membrane segmentation was the presence of background, as in cases of tightly packed cells. When tested for these conditions, the segmentation of the nuclear envelope was still possible. All the code and data are released openly through GitHub, Zenodo and EMPIAR.

## 1. Introduction

In 1951, cervical cells extracted from a patient called Henrietta Lacks at the Johns Hopkins Hospital were to become the first continuous cancer cell line [1]. The cells are widely known as *HeLa cells* (from *He*nrietta *La*cks) and have become a centrepiece of biomedical research, spanning from AIDS [2] to toxicity [3] to Zika [4] and, of course, cancer. As of 2021, PubMed contained more than 110,000 entries related to HeLa cells (https://pubmed.ncbi.nlm.nih.gov/?term=HeLa+[all+fields]). Since the cells were removed and kept without the patient’s knowledge or consent, which was not required at that time, many ethical and legal issues have also followed the HeLa cells [5–8].

The observation of the cells and their characteristics like shape, colours, or size and their relationship to health or disease is probably as old as the studies by van Leeuwenhoek and Hooke [9]. Whist the cell structure and its organelles have been well-known for many years, discoveries related to cell structure continue to appear in the scientific literature; searches in Google Scholar for terms such as *cell structure and function* (https://scholar.google.co.uk/scholar?as_ylo=2021&q=cell+structure+and+function) or *cell structure and transport* (https://scholar.google.co.uk/scholar?as_ylo=2021&q=cell+structure+and+transport) return more than 50,000 entries in the first four months of 2021 alone.

Today, sophisticated instruments, such as Electron Microscopes (EM), allow the observation with significant resolution, far greater than those of conventional light and fluorescence microscopes, and in turn allow the observation of smaller structures and provide great detail of the larger ones. Additionally, three-dimensional observation is possible by cutting very thin sections of a fixed sample with an ultra-microtome diamond knife [10] and acquiring images of the top face as each section is removed. The slice-acquisition process is known as *Serial blockface scanning EM* (SBF SEM) [11] and the output is illustrated in Figure 1(a) where selected images are positioned in three-dimensions according to their location in the volumetric stack. Figure 1(b) presents a zoom in to a single cell. One image is presented horizontally, and one orthogonal slice, or *orthoslice*, which is obtained from 300 images, is displayed vertically.

**Figure 1.**
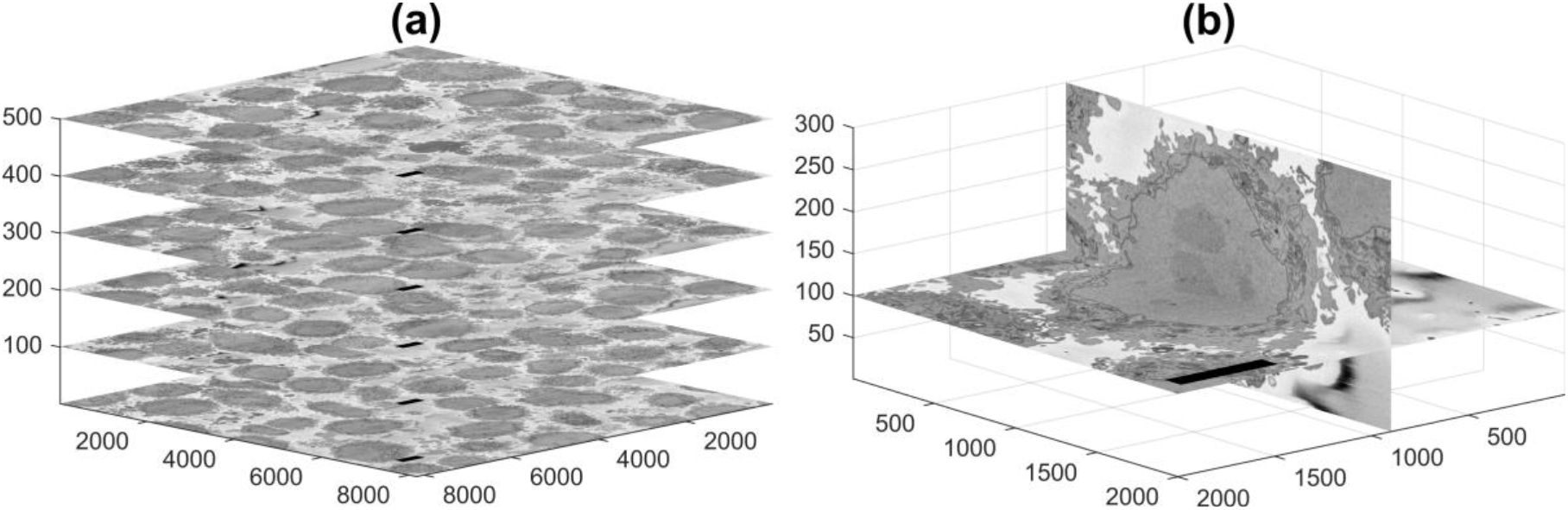
Representative slices of a 3D image stack acquired by Serial Block Face Scanning Electron Microscopy (SBF SEM) containing numerous HeLa cells. (a) Six of the 518 stack of electron microscopy images, and (b) For visualisation purposes two slices of a HeLa cell image have been presented vertical to each other. In both cases the units of the axes are in voxels and 5*μm* scale bars are shown towards the left of all horizontal images.

Cells are normally kept in shape by the plasma membrane, a phospholipid bilayer membrane that separates the internal aqueous environment of the cell and its organelles from the external environment [12]. In addition, the nucleus and chromosomes are surrounded by another bilayer membrane, the nuclear envelope [13]. The study of the cellular membranes has a long history and the study and discovery of cell structures “keep biologists glued to their microscopes” [14]. Cellular membrane receptors are important in conditions such as Alzheimer’s disease [15], cancer [16], Helicobacter pylori infection [17]. The geometry of the membranes is also important, for instance the shape [18,19], curvature [20,21] and protuberances [22] have been studied. The importance of the nuclear envelope in particular is related in processes such as viral infections [23], cancer [24], cardiovascular function [25] and has been an area of research for a long time [26,27]. Therefore, algorithms that provide accurate segmentation of the membranes of a cell are of great importance as the visualisation and analysis of the membranes and shape of cells could provide clues to understand the health or disease of cells and their organs [28–33]. The reader is referred to Lombard [34] for a historical review of the cell membranes.

Segmentation can be understood as the process of partitioning images or volumes into homogeneous non-overlapping regions [35,36], in its simplest case, one region is the background and the other region is an object or a foreground. Semantic segmentation identifies the pixels into a series of classes or labels, that have a particular meaning, like a person, a car, a cell or an organ [37]. Going further, instance segmentation is the process of detecting and segmenting each distinct object of interest appearing in an image [38]. For instance, if two cells are present in an image, semantic segmentation would identify them as cells, and instance segmentation would distinguish one cell from the other. Instance segmentation is more challenging than other pixel-level learning problems such as semantic segmentation, which deals with classifying each pixel of an image, given a set of classes. There, each pixel can belong to a set of predefined groups (or classes), whereas in instance segmentation the number of groups (instances) is unknown *a priori*.

Segmentation, identification and analysis of EM cellular images can be performed through manual processes [39–41], which can be distributed as *citizen science* where an *army of non-experts* [42,43] are recruited to provide non-expert human annotation, segmentation or classification through web-based interfaces (e.g. https://www.zooniverse.org/projects/h-spiers/etch-a-cell) [44]. Alternatively, computational approaches with traditional algorithms or deep learning approaches have been proposed to detect membrane neuronal and mitosis detection in breast cancer [45], mitochondria [46,47], synapses [48] and proteins [49]. Besides the well-known limitations of deep learning architectures, of significant computational power, large amount of training data and problems with unrelated datasets which show little value for unseen biological situations [50–55], the resolution of the EM data sets can enable or restrict their use for specific purposes. For instance, with voxel resolution of 10 nm and slice separation of 40 nm it is possible to observe well axons and dendrites [56], yet when voxel resolution is isotropic at 4 nm, an exquisite definition of macromolecular structures such as endoplasmic reticulum (ER) and microtubules are visible [57]. A notable contribution is CEM500K [58], a very large dataset of images, pre-trained models and curation pipeline for model building specific for Electron Microscopy. In addition, the specific nature of a semantic segmentation can allow traditional algorithms to provide satisfactory results, in some cases superior to deep learning approaches [59].

This paper describes an extension to previous work which focused on the segmentation the NE of a cell from a cropped volume [59,60]. In this work individual HeLa cells and their nuclei are instance segmented in 3D. The cells are identified and selected from a volumetric stack of 518 slices with8, 192 × 8, 192 pixels each. The number of cells to be identified and segmented can be selected as some cells will be close to the edges of the volume and will not appear complete. In order to segment an individual instance of one cell, volumes of 2, 000 × 2, 000 × 300 voxels, which contain a cell in the centre, are automatically cropped. Background and NE are automatically segmented and the resulting regions become the input to a series of steps of morphological distance, watershed, morphological operations that segment the cell from neighbouring cells. Thus the contributions of this work are: (a) the automatic identification and cropping of volumes that contain individual cells, and (b) the segmentation of the plasma membrane of a single cells and separating if from neighbouring cells. The rest of the manuscript is organised as follows:

All the code related to this work was performed in programming environment of Matlab^®^ (The Mathworks™, Natick, USA). Code, ground truth of one cell and EM images are available:

- *https://github.com/reyesaldasoro/Hela-Cell-Segmentation*.
- *https://doi.org/10.5281/zenodo.4590903*
- *http://dx.doi.org/10.6019/EMPIAR-10094*.

## 2. Materials and Methods

### 2.1. Cell preparation and Acquisition

HeLa cells were were prepared, embedded in Durcupan and observed with SBF SEM following the method of the National Centre for Microscopy and Imaging Research (NCMIR) [61]. SBF SEM data was collected using a 3View2XP (Gatan, Pleasanton, CA) attached to a Sigma VP SEM (Zeiss, Cambridge). In total, 518 images of 8, 192 × 8, 192 pixels were acquired. Voxel size was 10 × 10 × 50 nm with intensity [0 *−* 255]. Figure 1(a) shows six EM images positioned within the 3D stack. Initially, the data was acquired at higher bit-depth (32 bit or 16 bit) and after contrast/histogram adjustment it was reduced to 8 bit [60]. Images are openly accessible via the EMPIAR [62] public image database (*http://dx.doi.org/10.6019/EMPIAR-10094*).

### 2.2. Segmentation of background and identification of cells

One characteristic feature of the images is that the background, that is, the resin in which cells have been embedded, tends to be brighter than the cells, and is fairly uniform (Figure 2(a)). This allows the segmentation of the background on the basis of intensity. Cells are identified by a combination of intensity thresholding using Otsu’s algorithm [63], generation of super-pixels by detecting edges with a Canny edge detector [64] and morphological operators to clean the output (Figure 2(b)). It should be noted that an important limitation of the algorithm to identify cells is the presence of a brighter background. The background helps to identify individual cells and the proper delineation of the membrane, with its blebs and protrusions. When the presence of background is limited, as with cells that are close to each other, the segmentation will separate cells from each other, but the protrusions, which may belong to either cell may not be assigned as part of the cells. The presence of background is not a requirement for the nuclear envelope as it will be demonstrated below.

**Figure 2.**
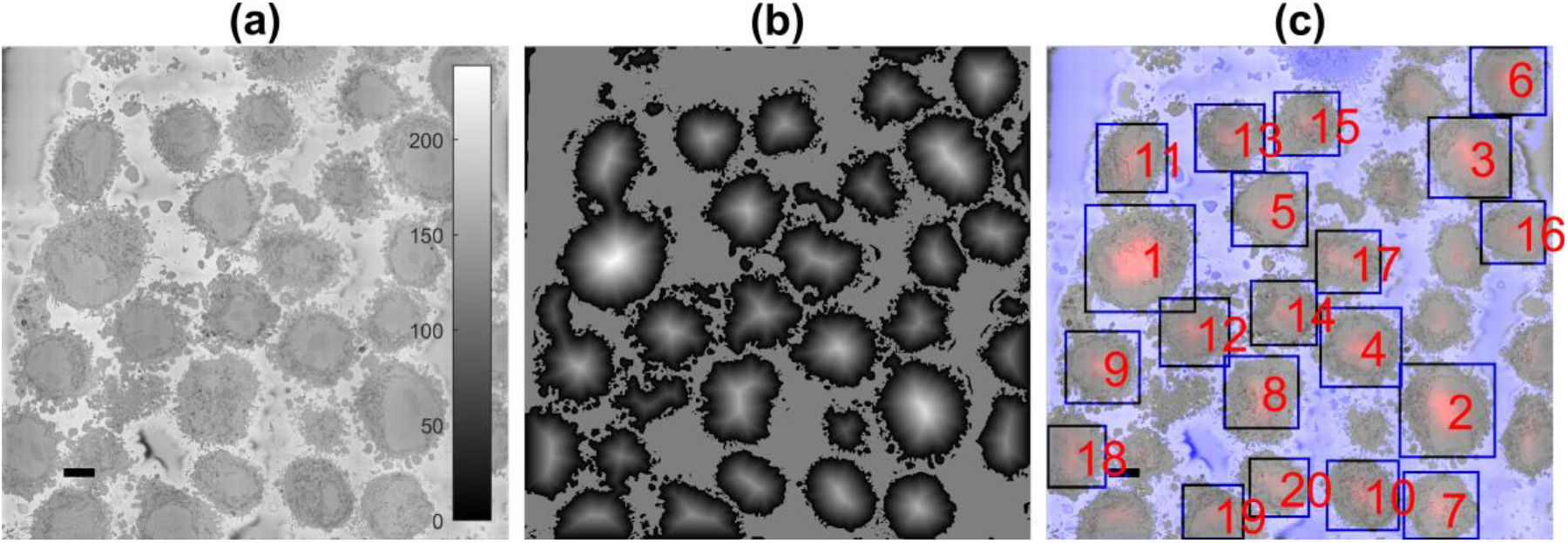
Automatic identification of cells from 8, 192 × 8, 192 images. (a) One representative slice with many HeLa cells. Scale bar corresponds to 5*μm*. (b) Illustration of the detected background (gray constant shade) and distance transform (gray to white) that corresponds to the cells, the larger the cell, the brighter the intensity of the transform. (c) Composite image of the slice as in (a), background as a purple shade and 20 detected cells, ranked in order of size. It should be noted that smaller cells are not selected as there was a limit of 20 in the present example.

The segmentation of the background leads to the identification of the cells through a calculation of a distance transform. Moreover, the distance allows to identify size of the cells as a larger cell will have pixels that further away from the background (Figure 2(b)). This could be understood as a topological analogy where the distance transform produces an *altitude* map and each cell corresponds to a *hill*. The ranking of the cells follows the altitude in descending order (Figure 2(c)). In some cases, it is possible that a single cell will provide more than a single peak, i.e., a range in the topological analogy. The process to identify these peaks as belonging to a single cell is to proceed iteratively from the highest peak and discard any other peak within a certain distance around it.

The number of cells to be identified can be pre-defined, e.g. 20 cells for the example of Figure 2(c). The identification is repeated for a number of slices of the 3D stack and the centroids of the cells are located in 3D. Then, these centroids are linked vertically to identify which of them correspond to the same cell as it should be noted that cell 1 will always be the largest cell of the particular slice and as the slices move up or down from what would be the equator of a cell, their size will change. Figure 3 illustrates the centroids located at every 20 slices (i.e., 26 slices were analysed) with the number from their corresponding slice and a coloured line indicating the centroids that were linked as a single cell and posteriorly cropped into to regions of interest of (2, 000 × 2, 000 × 300) voxels, in which one cell was centred. It should be noticed, that the only the largest 20 cells have been identified, with a few more smaller cells are still present. These cells may be small in the current slice as they are close to the edges, i.e. the poles, but could be larger in other slices.

**Figure 3.**
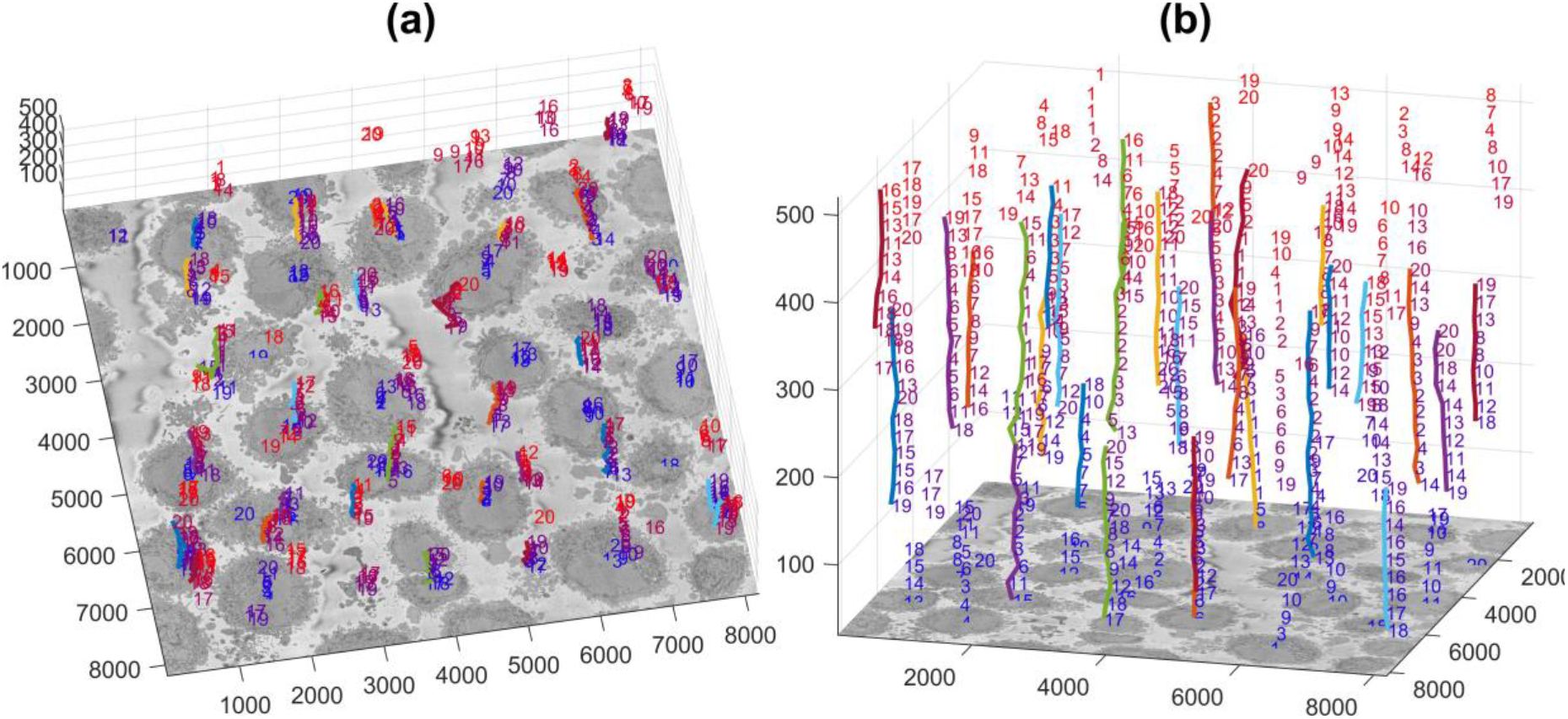
Centroids of cells that were identified per slice are displayed in three dimensions. Each number corresponds to the centroid of one cell that has been identified in a given slice. The numbers decrease according to the rank of the cell in that slice with 1 being the largest cell detected in that slice. The colour of the font varies from blue (lower slices) to red (higher slices) for visualisation purposes. A coloured line with a random colour is placed next to the centroids that were associated as a single cell. (a,b) show the same information from different points of view. The units of the axes are in voxels.

### 2.3. Semantic Segmentation of Nuclear Envelope

The process of segmentation continues now for each cell cropped within the 2, 000 × 2, 000 × 300 region of interest identified in the previous steps. The methodology for the automated segmentation algorithm of the Nuclear Envelope (NE) has been published before [60], but for completeness is summarised in this section.

In order to remove high frequency noise and therefore to improve segmentation, initially, all 518 HeLa cell EM images were low-pass filtered with a Gaussian kernel with size *h* = 7 and standard deviation *σ* = 2. Then, Canny edge detection [64] was used to determine the abrupt discontinuous in brightness or intensity changes between the NE and the neighbouring cytoplasm (outside the nucleus) and nucleoplasm (inside the nucleus).

The Canny edge detection resulted in some disjoint segments due to the NE variations in intensity and these segments were connected by dilation using a distance map from the edges. All pixels within an adaptive distance, i.e., 5 pixels, which grow depending on the standard deviation of the Canny edge detector, were connected as a single edge. Those pixels that were not considered as edges were labelled as a series of *superpixels*. Finally, several morphological operators were used to: remove regions in contact with the borders of the image, remove small regions, fill holes inside larger regions and close the jagged edges.

Starting from the central slice of the EM stack, e.g. the equator of the nucleus, the image-processing algorithm exploited the volumetric nature of the data by propagating the segmentation of the NE of one slice to the next, up and down from the centre. The NE of a previous slice was used to check the connectivity of disjoint regions or *islands* separate from the main nuclear region. The algorithm proceeded in both directions and propagated the region labelled as nucleus to decide if a disjoint nuclear region in the neighbouring slices was connected above or below the current slice of analysis. When a segmented nuclear region overlapped with the previous nuclear segmentations, it was maintained, when there was no overlap, it was discarded.

### 2.4. Semantic Segmentation of Cells

The segmentation of one cell from its neighbours is a relatively simple process when the background and nuclei have been previously identified (Figures 4(a,b)). In addition, since the current cell has been cropped into a region of interest where the cell nucleus is positioned near the centre of a 2, 000 × 2, 000 × 300 voxel volume, the segmentation becomes an instance segmentation as the cell will be identified as a cell and different from other cells that surround it.

**Figure 4.**
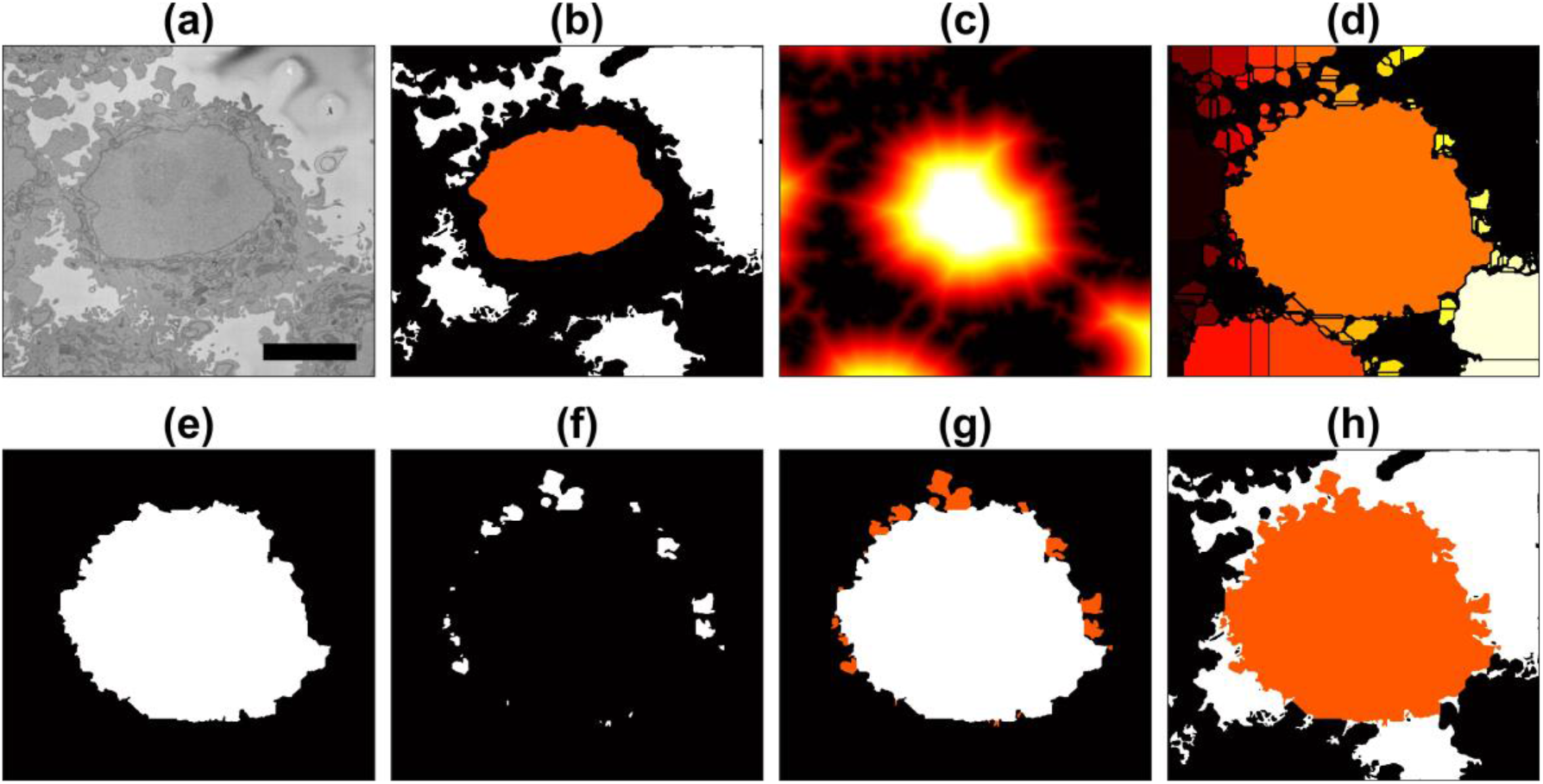
Illustration of the steps of the segmentation algorithm for a cell from neighbouring cells. (a) Region of interest (ROI) which contains one HeLa cell surrounded by background and other cells. The algorithm starts with the nuclear region and background. (c) Distance transform from the background. (d) Watershed transformation on the distance transform, all regions in the background are removed. (e) Central region from the watershed. (f) Small regions that are contiguous to the central region. (g) Addition of small regions, i.e., membrane protuberances. (h) Final result of the cell with the background in white and neighbouring cells in black. A 5*μm* scale bar is shown on panel (a).

The distance transformation from the background (Figure 4(c)) grows around regions with cells and since there will be one larger *hill* at the centre, the distance transformation can be segmented with a watershed algorithm [65]. The watershed is useful to separate one cell from other cells within the field of view. Watersheds are well-known for over segmenting and other artefacts so the central and largest region is selected as the cell (Figure 4(d,e)). This region is morphologically *opened* with large structural elements to remove protruding artefactual regions product of the watershed (Figure 4(e)). This returns a fairly round cell which will not include the natural protrusions (i.e., pseudopods or cell membrane extensions) of the cell. Thus, regions that are contiguous to this central region and surrounded by background are identified and merged with the cell (Figure 4(f,g)). The final segmentation of thecell with the inclusion of these are natural protrusions or protuberances of the cell membrane can be observed in Figure 4(h).

### 2.5. Quantitative Comparison

In order to assess the accuracy of the segmentation algorithm, two different pixel-based metrics were used: accuracy (AC) and Jaccard similarity index (JI) [66]. Both metrics arise from the allocation of classes (Nucleus, Cell, Background) to every pixel of an image and the correct or incorrect prediction of the class byt the segmentation algorithm. For each pixel, four cases exist: true positive (TP), which correspond to pixels which were correctly predicted as a certain class (e.g. nucleus), true negative (TN), false positive (FP), and false negative (FN). Thus, accuracy is calculated as:

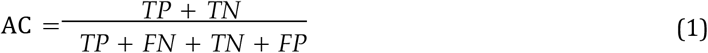

and Jaccard index is calculated as:

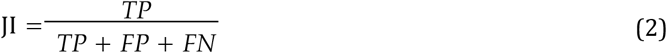

It should be noticed that JI is a more rigorous metric since TN, or correctly segmented background pixels are not taken into account. A very small region of interest surrounded by a very large background could produce a very large accurate result due to the presence of TN.

The AC and JI metrics were calculated for three different scenarios: (a) the segmentation of the cell without the nucleus, (b) the cell including the nucleus and (c) the nucleus only. Figure 5 illustrates these cases for a sample slice of the data.

**Figure 5.**
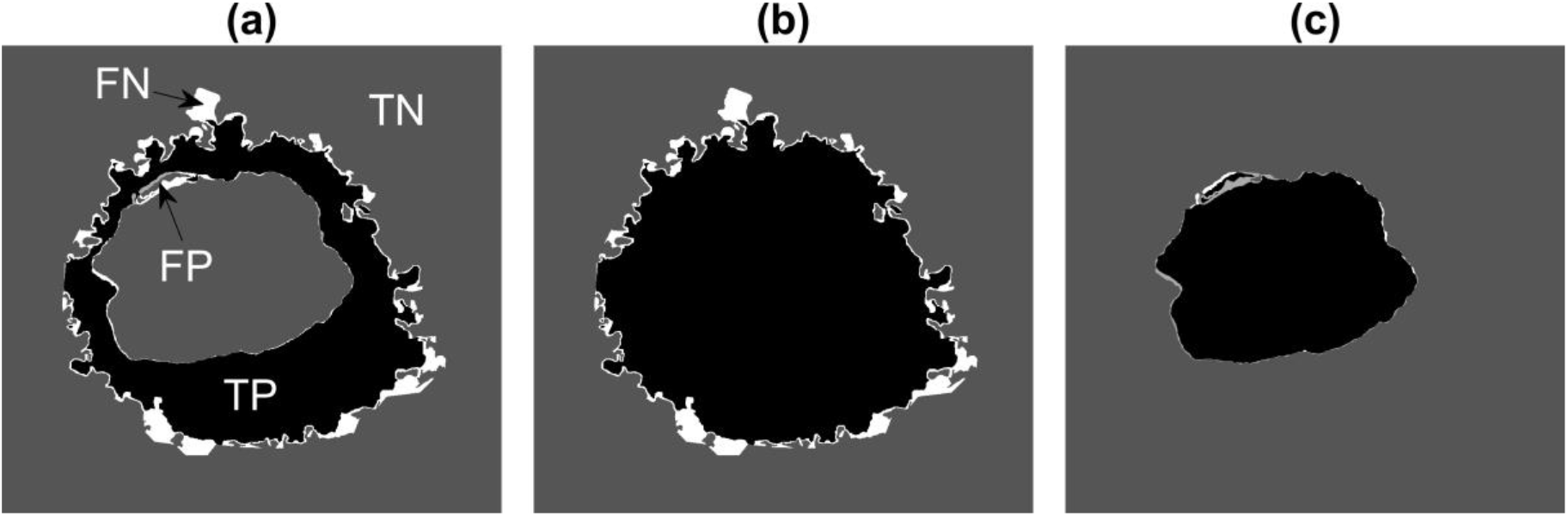
Illustration of the pixel-based metrics. True Positives (TP, black), true negatives (TN, dark gray), false positives (FP, light gray) and false negatives (FN, white) are presented with increasing gray level intensity. (a) Cellular region excluding nucleus. (b) Entire cellular region. (c) Nucleus. FN are far more common than FP in (a,b) as some convoluted regions of the cell have not been segmented.

## 3. Results and Discussion

A volume of interest of 8, 192 × 8, 192 × 518 voxels was analysed as previously described. The cell identification process was run for 26 slices, i.e., every 20 slices. At each slice, the algorithm was set to detect 20 cells. When the results of all slices were linked, these identified 30 cells in total. The only manual intervention of this process is to select the number of cells per slice and the number of slices to be analysed. Thirty cells automatically identified is comparable than those analysed through citizen science approaches, i.e., 18 [44].

These cells were automatically cropped into invididual ROIs and saved in separate folders as 300 2, 000 × 2, 000 Tiff images. The ROIs are illustrated in Figure 6. It was observed that some of the ROIs were close to the top or bottom (e.g. ROIs 1, 2, 29, 30) and thus would contain only part of a cell and the rest would be outside the field of view in the vertical direction. Similarly, the centroids of other ROIs were close to the edges of the volume and thus are not centred but rather positioned towards a side of the ROI (e.g. 6, 14). Since the actual centroid is recorded, it is possible to determine how far away (i.e., absolute distance) are the centroids from the edges of the initial 8, 192 × 8, 192 × 518 volume. The centroid of ROIs 6, 14, 15, 27 and 28 were less than 500 pixels from the edge. Considering that these ROIs are 2,000 pixels wide, this indicates that the centroid is considerably distant from the centre, closer to the edges than to the centre of the 2, 000 × 2, 000 × 300 volume. It should be noted that one assumption of the nuclei segmentation is that the nuclei are centred in the ROI. In addition, by being located away centre allows the possibility of two partial cells occurring in the same ROI.Thus these six ROIs were discarded and not further processed. This decision was taken manually but it could be automated by determining a certain margin from the edges. Alternatively, it could be possible that instead of fixing the size of the ROI to be 2, 000 × 2, 000 × 300, a different size could be allocated so that the centroid of the cell is always at the centre of the volume.

**Figure 6.**
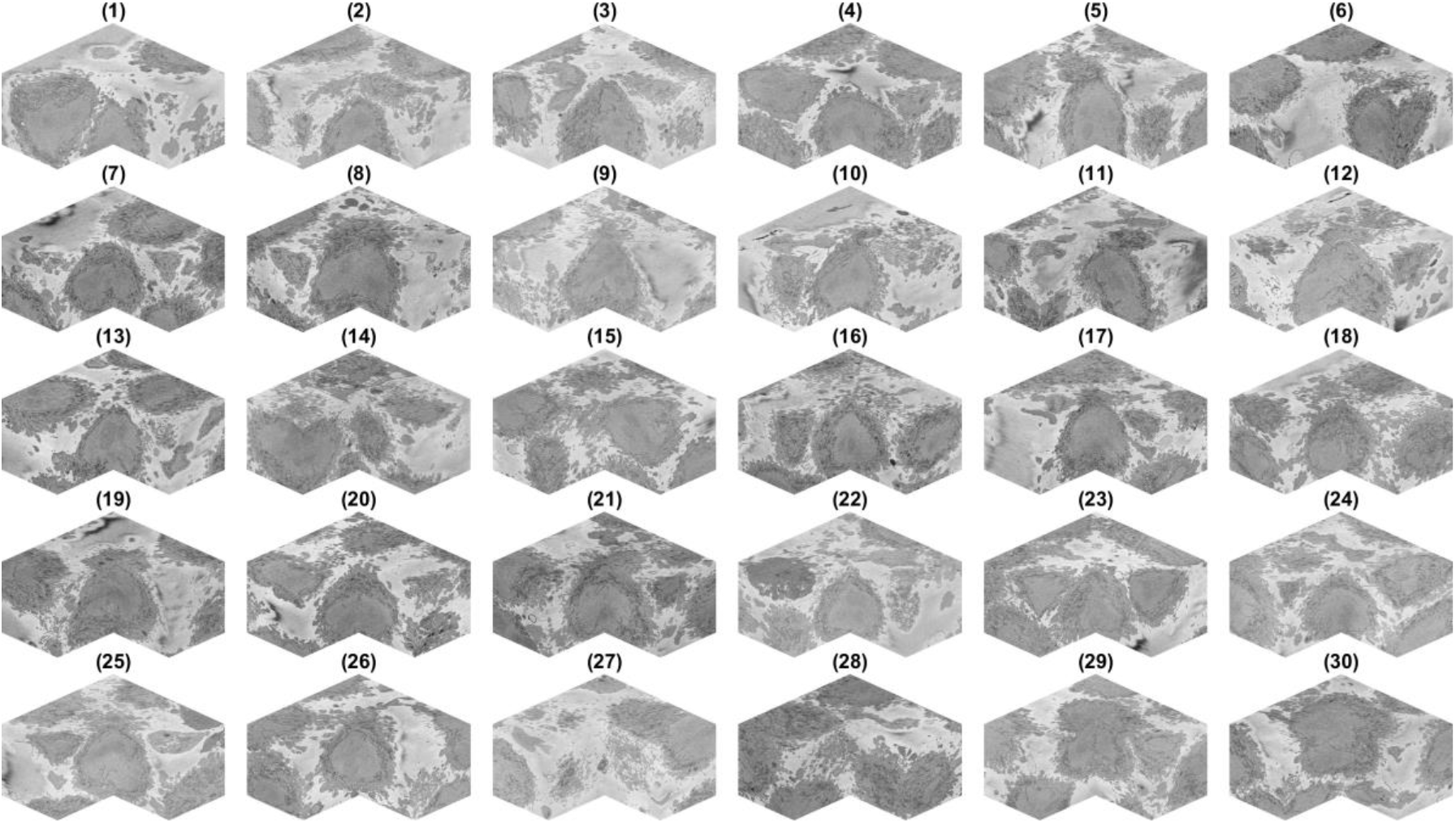
Regions of interest (ROI) cropped from the volume. Thirty regions of interest were detected and cropped. For each region of interest a corner is removed to show the cell, which should be centred. It should be noted that some cells are not in the centre but rather positioned towards the bottom of the volume (e.g. 1), the top (e.g. 30) or the sides (e.g. 6, 14, 15, 27, 28).

The situation of cells that are close to the top (29, 30) and bottom (1, 2) is slightly different. Whilst the cell is not centred *vertically* it is still centred *horizontally* within the plane of the images and it can be segmented successfully if the central slice to be processed is adjusted with the centroid location. The segmentation will not be of a complete cell but only part of it, so the segmentation may appear as a *hat* when it is in the lower section of the volume or a *bowl* if it is in the upper section. If metrics like volume are to be obtained, then these cases are not to be used. In this manuscript we are interested in the observation, rather than a quantification. These cells were thus not discarded from the analysis.

The cells and nuclei in the remaining 25 ROIS were segmented and are displayed separately in Figure 7 and together in Figure 8 with transparent membranes and solid NEs. Colours have been assigned randomly in Figure 8 for visualisation purposes together with one slice of 8, 192 × 8, 192 pixels to give reference. The positions of Figure 6 are maintained in Figure 7 to facilitate the comparisonbetween the ROIs and the segmentations. Whilst at this resolution it is not easy to observe details,it can be seen that the 25 cells and nuclei were successfully segmented. Geometric features fromthese surfaces could be extracted if different populations of cells, for instance treated/untreated, wild type/mutant, healthy/infected were to be compared statistically. We do not conduct this analysis as all these cells correspond to a single experimental condition.

**Figure 7.**
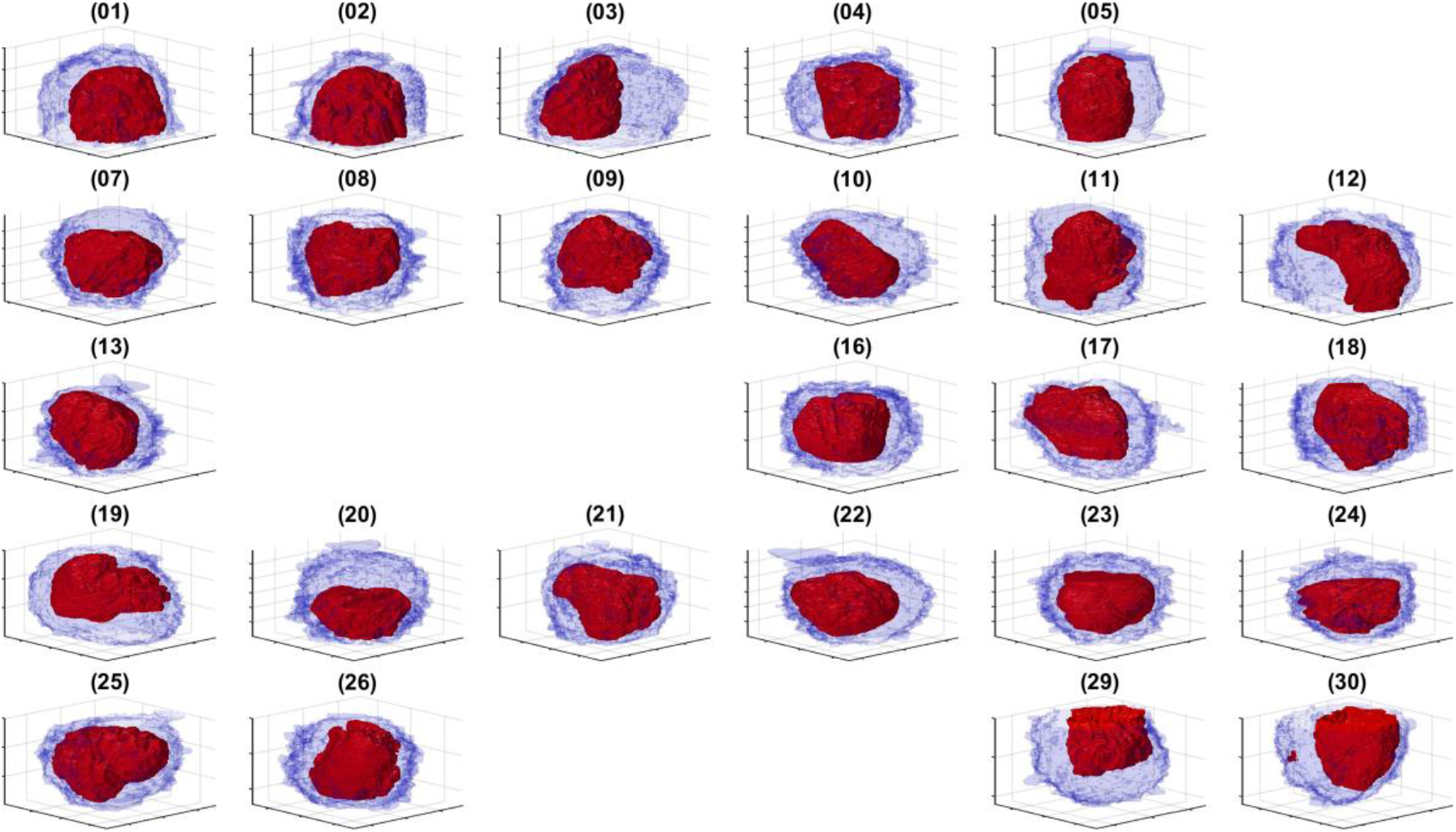
Rendering of the cell and nuclear envelope (NE) of 25 cells. For each case, the NE is rendered in red without transparency and the cell membrane is rendered in blue with transparency. The cells in ROIs 6, 14, 15, 27, 28 were located on the edges of the volume and the centroids were too close to the edges and thus discarded. For comparison purposes the cells are placed in the same locations as in Figure 6, and the region of interests (ROIs) that were discarded are blank.

**Figure 8.**
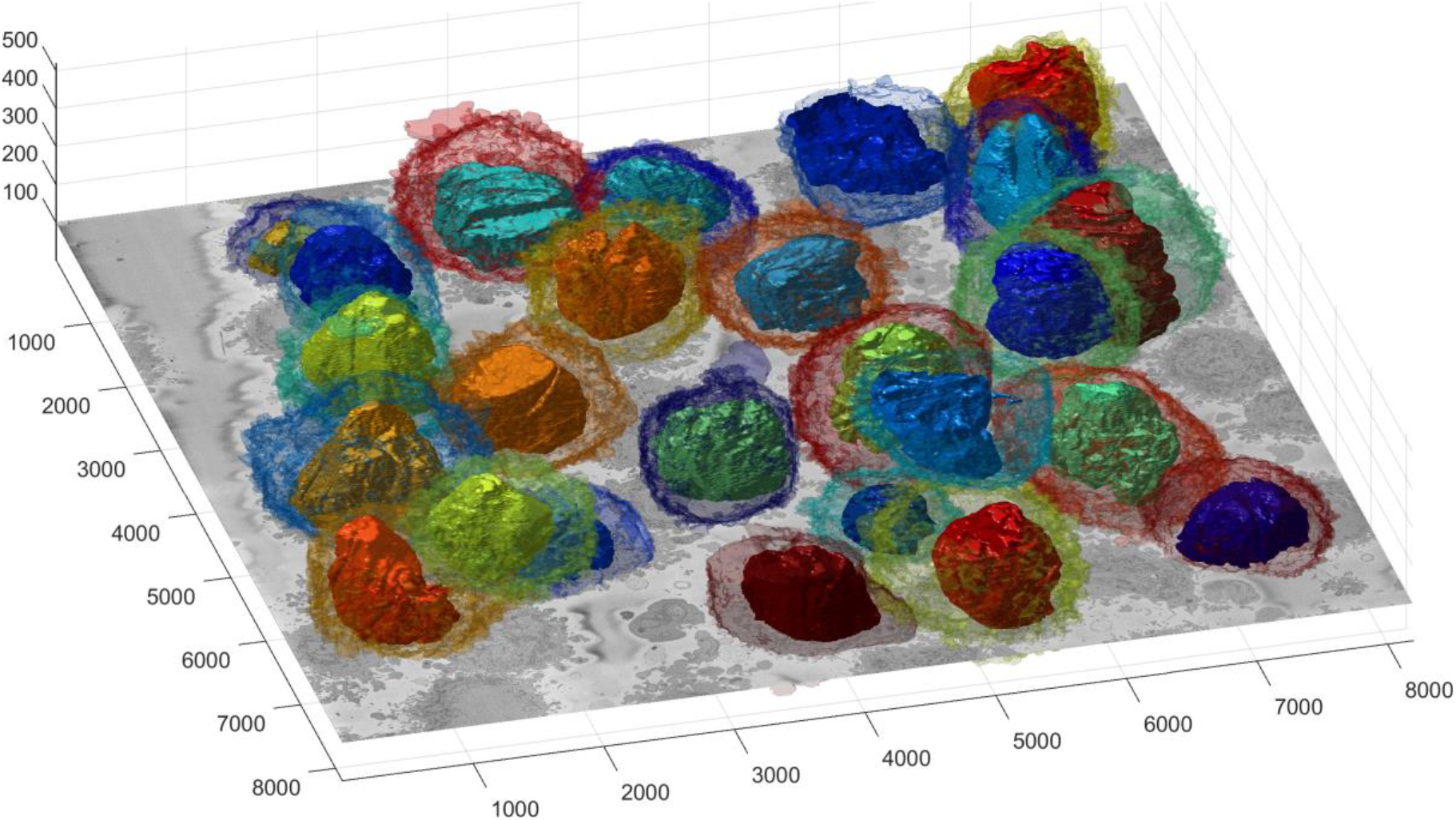
Illustration of segmentation of 25 cells and nuclear envelopes (NE). The cells were segmented from a 8, 192 × 8, 192 × 518 voxel region. Slice number 100 out of 518 is displayed for context. All nuclei are shown solid and all cell membranes are shown in transparency, colours have been assigned randomly for visualisation purposes. The units of the axes are in voxels.

To better observe the details, the rendering of four cells is shown in Figure 9. Left and central columns show the cell membrane rendered with high transparency and the NE rendered as a solid surface from two different points of view. Right column shows the cell membrane without transparencyfrom the same point of view as the central column. The first interesting observation is that the nuclei seems to be far more different than the cells themselves in terms of the smoothness or ruggedness ofthe surface and the distribution of the nucleus within the cell. The first cell (a,b), rendered in red is relatively smooth whist the other cells have far more grooves on the surface. It is also interesting to observe that the nucleus does not appear to be in the same position for these cells, leaving space in the top (b), bottom (h,k) and roughly even (e). These characteristics, the uneven distribution of the nuclei and their variability as compared with the cells, are also visible in Figure 11, and its magnified version in Figure 10, which show the results overlaid on four slices of the data with nuclei with a green shade and cells with a red shade. Numbers to identify the ROIs have been added, but it should be noted that some cells are not visible within certain slices and may be positioned over other cells not included in the analysis. Some cells (e.g., 3, 8, 12, 19) appear to be polarised to a side with the perinuclear region with numerous organelles visible. The results shown in Figure 10 illustrate well boundaries between several cells and the complexity of the plasma membrane can be appreciated..

**Figure 9.**
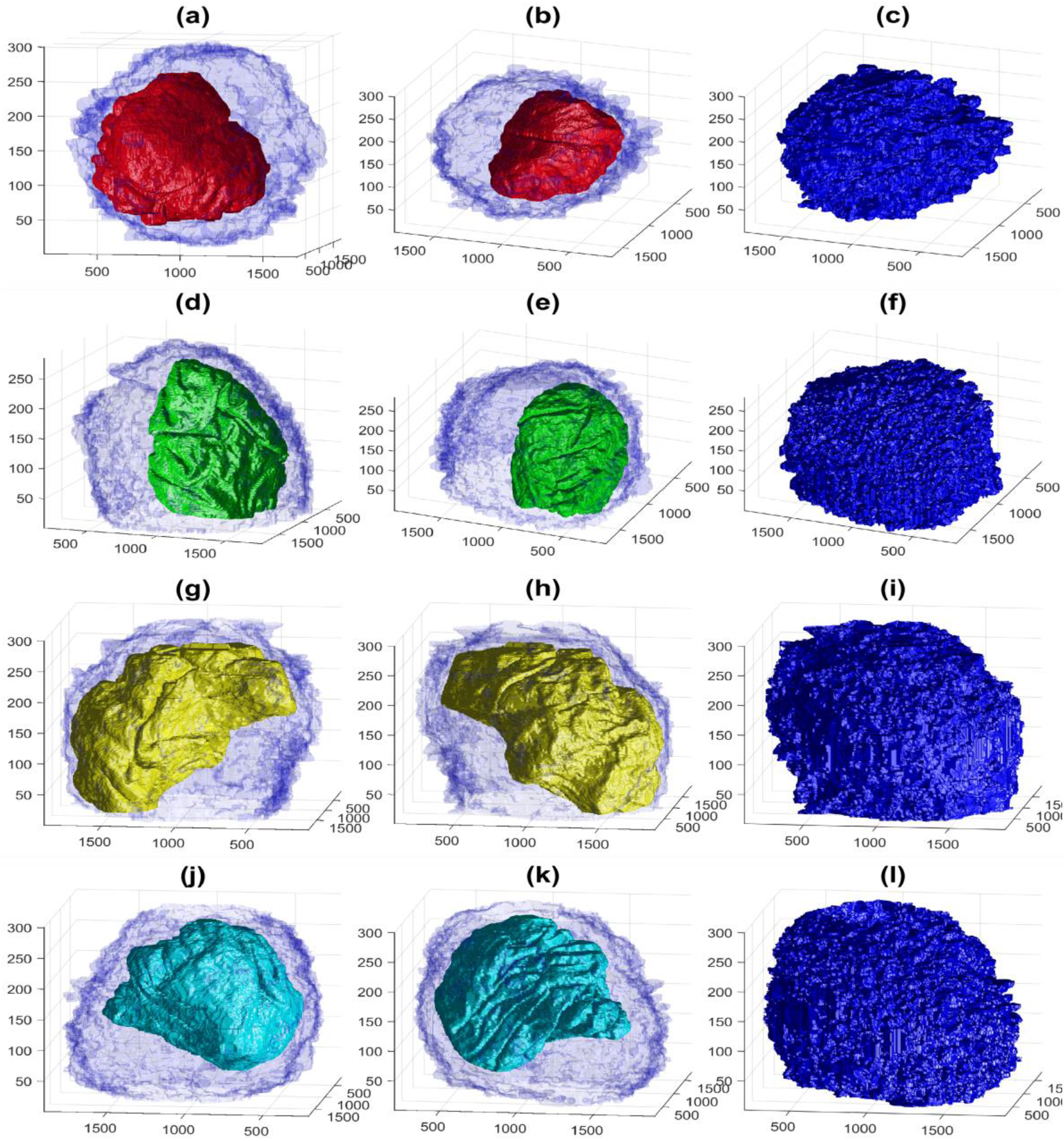
Four examples of volumetric reconstruction of the nuclear envelope (NE) and the cell membrane of HeLa cells. In all cases, each row corresponds to a single cell observed from different view points. Left and centre columns show the cell membrane with transparency. Right column the cell membrane without transparency from the same view point as centre column. The volume of interest is 2, 000 × 2, 000 × 300 voxels and the units of the axes are in voxels. (a,b,c) Region of Interest (ROI) 23, NE is shown in red and cell in blue. Notice the relative smoothness except for one groove along the cell and the concentration on the lower part of the cell. (d,e,f) ROI 3, NE is shown in green, notice the ruggedness of the NE with numerous grooves and the concentration of the nucleus towards one side of the cell. (g,h,i) ROI 12, NE is shown in yellow. Notice the distribution of the nucleus concentrated on the upper part of the cell. (j,k,l) ROI 19, NE is shown in cyan. The surface of the NEs appears more distinctive than those of the cells.

**Figure 10.**
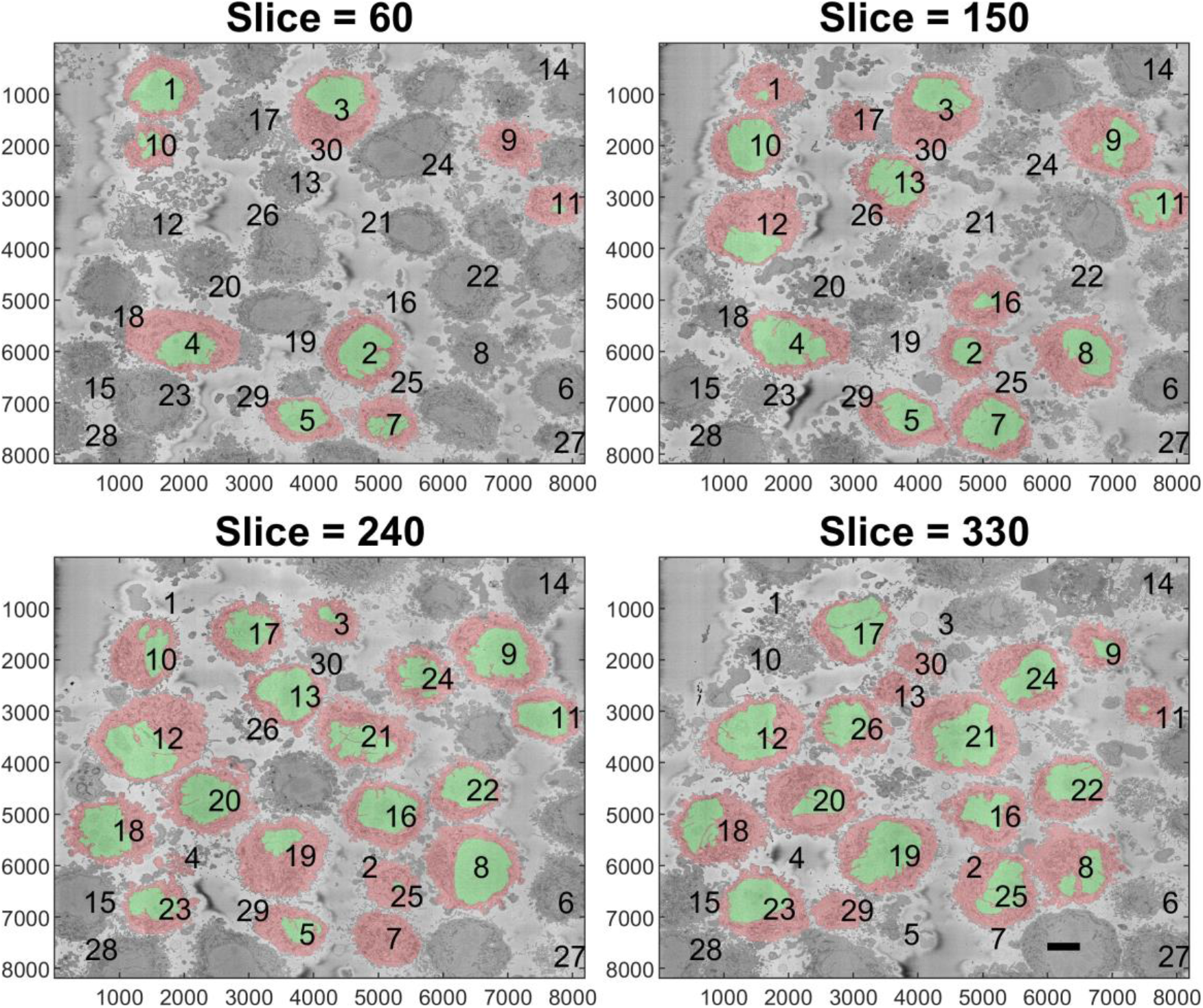
Final illustration of the results: three slices with the segmentations overlaid. In each slice, the cells are highlighted with a red shade and the nuclei are highlighted with a green shade. Numbers are added to aid the localisation of the particular cell. Notice that some of the numbers correspond to cells that are not visible in that particular slice. The units of the axes are in pixels and a scale bar indicating 5*μm* is shown in slice 330.

**Figure 11.**
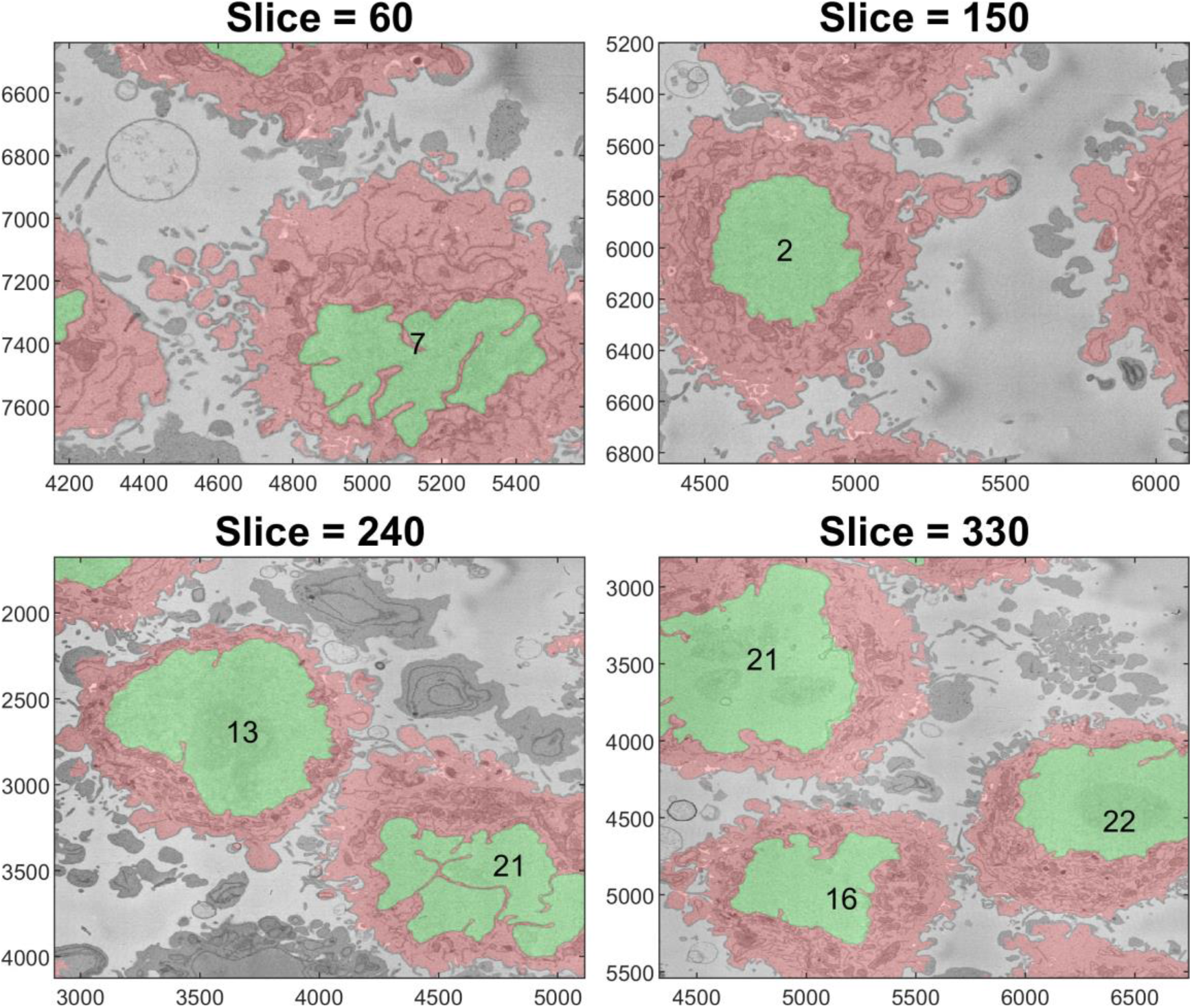
Final illustration of the results: three slices with the segmentations overlaid. The units of the axes are in pixels (a).

One final observation of Figure 11 is that there is one cell that was not identified by the algorithm; the cell in the centre of slice 240 in between cells 13, 21, 16, 19 and 21. This may be due to two factors, first, the algorithm was selected to identify only 20 cells per slice. Slice 240 shows 19 cells, and it should be remembered that cells 6, 14, 15, 27 and 28 were discarded. Thus it may be that by using a larger number per slice, e.g., 30 this cell would be selected. Second, that cell in particular is rather *flat* in the vertical dimension if compared with cells 13 and 16 that are visible in slices 150, 240 and 330. Since the algorithm ranks cells by their size, the cell may have been smaller in size and thus not considered. Again, by extending the number of cells per slice this could include this cell and others in the analysis.

The results for the cell for which ground truth for the nuclear envelope and cell membrane was available were the following. For the cell not including the nucleus *AC* = 0.9629, *J I* = 0.8094, for the cell and the nucleus *AC* = 0.9655, *J I* = 0.8711 and for the nucleus alone *AC* = 0.9975, *J I* = 0.9665. The algorithm to segment the nucleus provided excellent results, and it had previously been reported that it outperformed several deep learning architectures [59]. The small differences between the segmented nucleus and that of a manual expert segmentation are due mainly to the calculations of the thickness of the NE and small invaginations (Figures 12, 13 right column). As it would be expected due to the complexity of the cell membrane, the values for the cell are relatively lower than those of the NE. Figures 12, 13 show that there are relatively few FP in comparison to the FN. This is a characteristic of the segmentation algorithm, i.e., it is *cautious* to include the protuberances that surround the cell. These regions are better observed in Figure 13, which zooms in and illustrate regions where these FN appear. It can be perceived that in some cases, it is difficult to decide if a small region belongs to the cell of interest or to the neighbouring cells. It is well known that it is difficult to establish the *truth* and there could be significant inter- and intra-observer variability [44].

**Figure 12.**
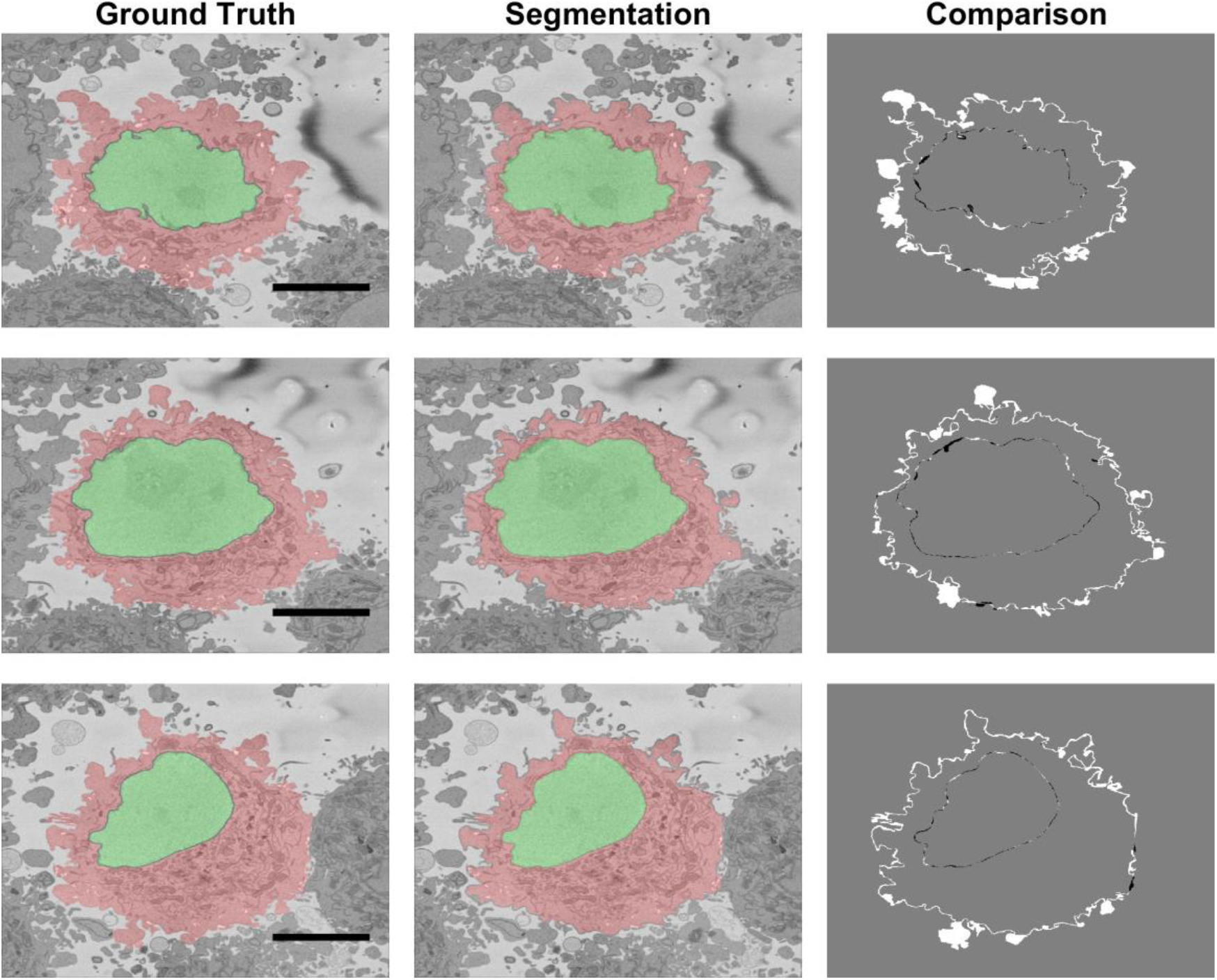
Comparison between ground truth (GT) and segmentation result obtained from the segmentation algorithm shown in three slices of the stack. Left column illustrates the GT with shades of green for the nucleus and shade of red for the cell. Centre column shows the result of the segmentation algorithm. Right column show the comparison between GT and the results with FN in white, FP in black and both TP and TN in gray. Large white regions correspond to distinction between neighbouring cells. A 5*μm* scale bar is shown in the GTs.

**Figure 13.**
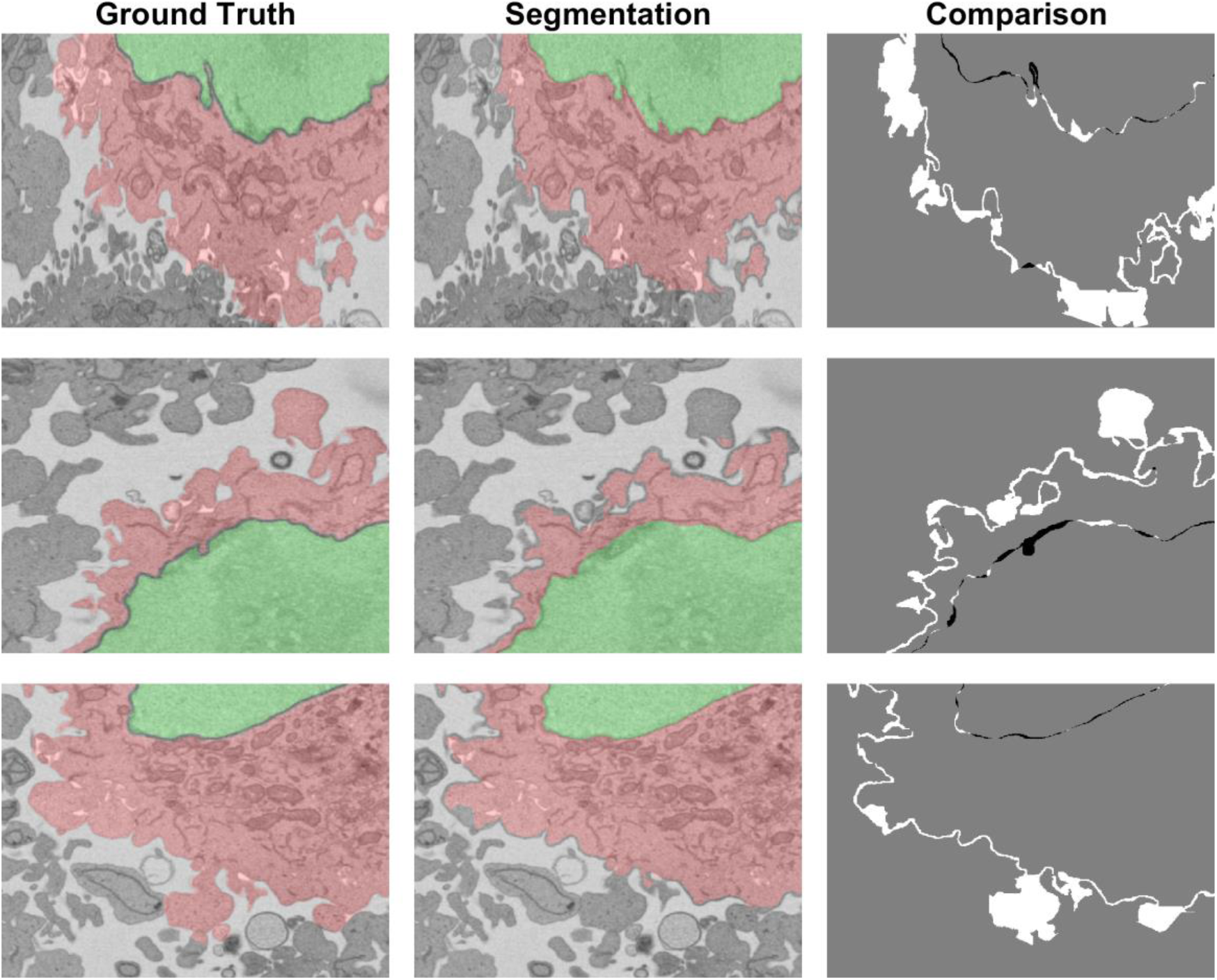
Magnified version of Figure 12. The white regions of the comparison correspond to the FN, i.e., regions that were not selected by the algorithm. It can be observed that some of these regions, which could belong to either of the neighbouring cells, would be difficult to identify to a human expert.

This situation is further illustrated in Figure 14, which shows several slices of the volumetric stack. Asterisks have been placed next to regions which could belong to either the cell of interest or neighbouring cells. In all the cases illustrated, the algorithm did not include these as part of the cell. The risk of modifying the algorithm to include the protuberances that were not included would be that the cell could grow into neighbouring cells. Protuberances surrounded mostly by background and not so close to other cells are better segmented as indicated by black diamonds. If the cell surfaces were to be analysed for counting filipodia or to distinguish filipodia from lamellipodia, the algorithm would have to be refined.

**Figure 14.**
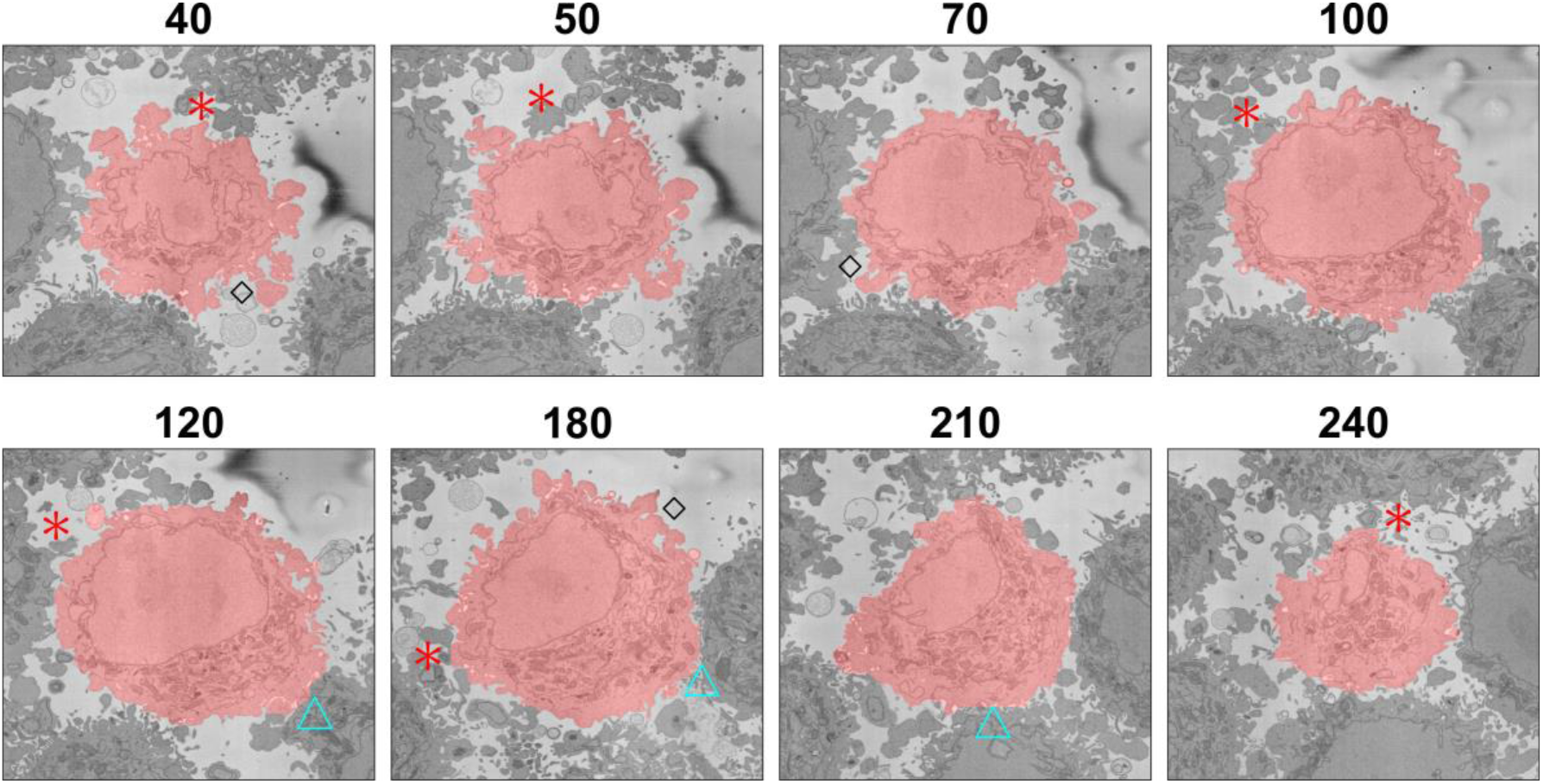
Illustration of segmentation at several slices of one cell. The segmentation is indicated with a red shade over the cell. Red asterisks indicate regions where the segmentation did not include protuberances that could belong either to the cell or to neighbouring cells. Blue triangles indicate regions where two cells are close together and the segmentation tends to a straight line between the cells. Black diamonds indicate convoluted protuberances that have been correctly segmented. The slice number relative to the stack of 300 is indicated above each slice.

The segmentation of one cell from other neighbouring cells requires the presence of a certain amount of background surrounding the cell, as this is used to form the distance map, which will later be segmented with the watershed. When two cells are very close together and there is no background, or very little of it, in that region the segmentation tends to be a straight line that bisects the cell with its neighbouring cells. This is illustrated in Figure 14 with cyan triangles next to those boundaries. As mentioned earlier, false negatives, i.e., the convoluted regions not included, outnumber false positives; false negatives constituted 3.46%of voxels whilst false positives were only 0.25%for the cell without nucleus and 3.31%and 0.14%for the cell including the nucleus. For reference the true positives were 15.7/23.33%and the true negatives 80.5/73.2%. Thus, the algorithm is a lower-bound estimation of the cell and is not including as part of the cell regions that could belong to either of two neigbouring cells, around 3%for the one cell for with GT was available. Again, if a very precise cell-cell interaction based on the surfaces were of interest, the algorithm would have to be refined. Still for many other applications such as measurements of volumes, position, polarisation, this algorithm provides a very good approximation of the cell membrane and an excellent segmentation of the nuclear envelope. Another application where this algorithm can be used is for counting mitochondria or other structures on a *per cell basis*, so that each instance of a mitochondria can be allocated to the correct cell.

As an indication of the computational complexity, the times to run the following processes were recorded in an Alienware m15 R3 Laptop with an Intel®Core™i9-10980HK CPU, 2.40 GHz with 32 GB RAM and running MATLAB®(Mathworks™, Natick, USA) Version: 9.8.0.1417392 (R2020a). (1) Detection of 20 cells in 26 slices and joining to identify 30 cells, **48 s**, (2) Cropping 30 ROIs and saving 300 Tiff images in 30 folders **9.1 min** (18.2 sec/ROI), (3) Segmentation of one nuclei **14.9 min** (2.98 sec/slice, (4) Segmentation of one cell **13.2 min** (2.64 sec/slice). These times are significantly faster than the manual segmentation that can take around 30 hours to segment one nuclear envelope alone [60].

So far, one of the strongest limitations of the algorithm is the presence of a bright background. To further test the segmentation algorithms, we identified cells with different settings that the 8, 192 × 8, 192 × 518 volume previously described. Two publicly available EM datasets from the **Cell Image Library - (CIL)** (*http://cellimagelibrary.org/images/50051* and *http://cellimagelibrary.org/images/50061*) were analysed. These sets consisted of Chlamydia trachomatis-infected HeLa Cells also embeded in Durcupan and acquired with SBF SEM using a Gatan automated 3View system (Gatan Inc.) [67]. The intensities were remarkably different from the previous data set as illustrated in Figures 15(a,d).

**Figure 15.**
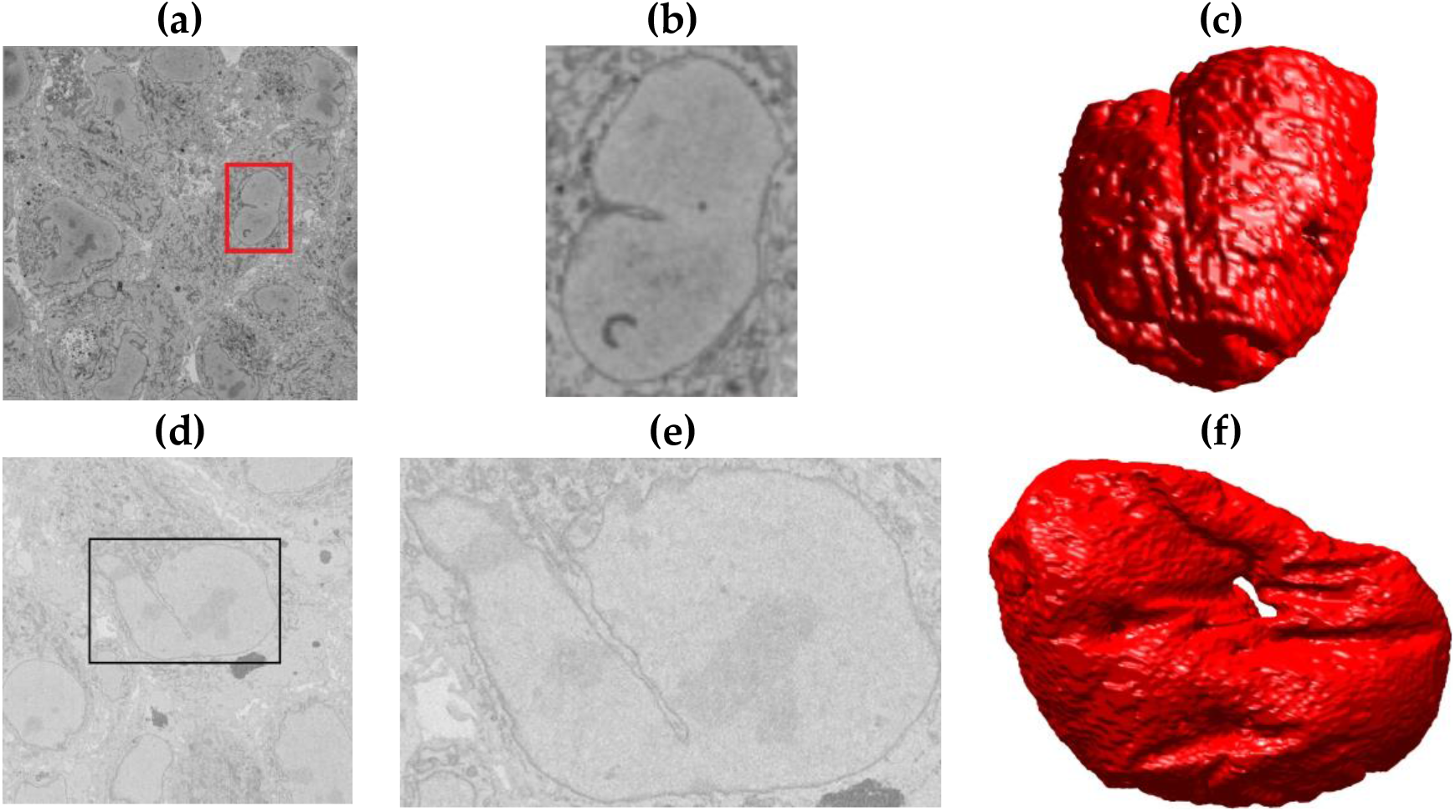
Illustration of the Serial Block Face Scanning Electron Microscope (SBF SEM) images containing monolayers of Chlamydia trachomatis-infected HeLa cells. (a) A representative image from the Cell Image Library *CIL*50051 data set. The volume has 3200 × 3200 × 413 voxels and voxel size is 3.6 × 3.6 × 60 nm. (b) A region of interest (ROI) with one nucleus, which corresponds to the red box in (a). (c) Rendering of the nuclear envelope (NE) of this cell. (d) One representative image from the Cell Image Library *CIL*50061 data set. The set has 2435 × 2489 × 406 voxels and voxel size 8.6 × 8.6 × 60 nm. (e) ROI with one nucleus corresponding to the black box in (d). (f) Rendering of the NE of this cell.

Notice especially the low contrast of (d). It was observed that the closeness between cells resulted in a very limited background and thus the analysis was restricted to the NE from manually cropped ROIs (Figures 15(b,e)). The nuclei were segmented and results were considered very good through a visual inspection (Figures 15(c,f)).

One further observation was relevant. The surface on Figure 15(f) showed a hole so it was interesting to display the surface of the NE in a different way with the NE rendered as a mesh with no face colour, edges in black and with transparency (Figure16). Axis are added for reference. Besides the evident hole, the NE has other deep crevices that nearly connect two opposite sides of the NE as can be observed in the lower part of (b).

**Figure 16.**
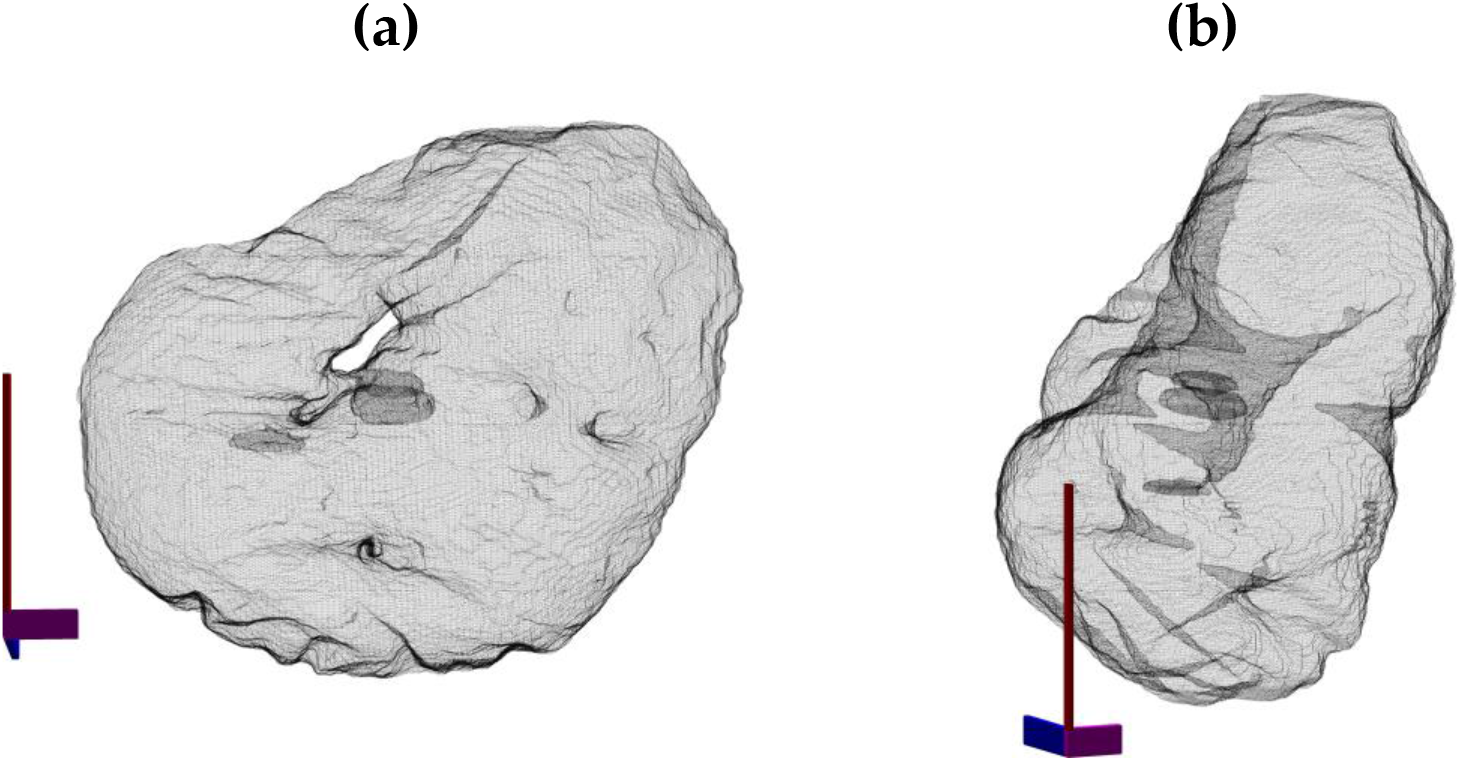
Illustration of the nuclear envelope from the data set CIL50051. The surface is displayed as a mesh with transparency to show the hole of the nuclear envelope (a) and the crevices that go deep inside the nucleus. Notice in (b) how these invaginations nearly connect separate sides of the NE.

In this paper, an algorithm to segment instances of HeLa cells as observed with electron microscopy was described. The algorithm can be run automatically and only requires some parameters (number of cell per slice, number of slices, cells to be discarded) were set manually. The results of the algorithm were very accurate for the segmentation of the nuclear envelope and lower for the cell membrane, as it would be expected due to the significantly more complex geometry of the latter. The algorithm has some limitations that have been discussed above, most important the presence of a bright background and the cautious delineation of the membrane where some regions that could belong to either a cell or its neighbour were not included in either. For a general analysis of the cell surface, the algorithm provides good results, and if details of the protrusions of the cell were required, further work could consider a post-analysis of all regions not included once all cells have been segmented. Future work will be dedicated to the segmentation of other organelles such as mitochondria and Golgi apparatus.

## References

1. Masters, J.R. HeLa cells 50 years on: the good, the bad and the ugly. Nature Reviews Cancer 2002, 2, 315–319. doi:10.1038/nrc775.

2. Chesebro, B.; Wehrly, K.; Metcalf, J.; Griffin, D.E. Use of a new CD4-positive HeLa cell clone for direct quantitation of infectious human immunodeficiency virus from blood cells of AIDS patients. The Journal of Infectious Diseases 1991, 163, 64–70. doi:10.1093/infdis/163.1.64.

3. Bich-Loan, N.T.; Kien, K.T.; Thanh, N.L.; Kim-Thanh, N.T.; Huy, N.Q.; The-Hai, P.; Muller, M.; Nachtergael, A.; Duez, P.; Thang, N.D. Toxicity and Anti-Proliferative Properties of Anisomeles indica Ethanol Extract on Cervical Cancer HeLa Cells and Zebrafish Embryos. Life (Basel, Switzerland) 2021, 11. doi:10.3390/life11030257.

4. Li, L.; Collins, N.D.; Widen, S.G.; Davis, E.H.; Kaiser, J.A.; White, M.M.; Greenberg, M.B.; Barrett, A.D.T.; Bourne, N.; Sarathy, V.V. Attenuation of Zika Virus by Passage in Human HeLa Cells. Vaccines 2019, 7. doi:10.3390/vaccines7030093.

5. Kemet, S. Insight Medicine Lacks - The Continuing Relevance of Henrietta Lacks. The New England Journal of Medicine 2019, 381, 800–801. doi:10.1056/NEJMp1905346.

6. Witze, A. Wealthy funder pays reparations for use of HeLa cells. Nature 2020, 587, 20–21. doi:10.1038/d41586-020-03042-5.

7. Wolinetz, C.D.; Collins, F.S. Recognition of Research Participants’ Need for Autonomy: Remembering the Legacy of Henrietta Lacks. JAMA 2020, 324, 1027–1028. doi:10.1001/jama.2020.15936.

8. Beskow, L.M. Lessons from HeLa Cells: The Ethics and Policy of Biospecimens. Annual review of genomics and human genetics 2016, 17, 395–417. doi:10.1146/annurev-genom-083115-022536.

9. Ribatti, D. An historical note on the cell theory. Experimental Cell Research 2018, 364, 1–4. doi:10.1016/j.yexcr.2018.01.038.

10. Peddie, C.; Collinson, L. Exploring the third dimension: Volume electron microscopy comes of age. Micron 2014, 61, 9–19. doi:10.1016/j.micron.2014.01.009.

11. Denk, W.and Horstmann, H. Serial block-face scanning electron microscopy to reconstruct three-dimensional tissue nanostructure. PLoS Biol. 2004, 2.

12. Nagle, J.F.; Tristram-Nagle, S. Structure of lipid bilayers. Biochimica et biophysica acta 2000, 1469, 159–195.

13. De Magistris, P.; Antonin, W. The Dynamic Nature of the Nuclear Envelope. Current biology: CB 2018, 28, R487–R497. doi:10.1016/j.cub.2018.01.073.

14. Kwok, R. Cell biology: The new cell anatomy. Nature News 2011, 480, 26. doi:10.1038/480026a.

15. Amadoruge, P.C.; Barnham, K.J. Alzheimer’s disease and metals: a review of the involvement of cellular membrane receptors in metallosignalling. International Journal of Alzheimer’s Disease 2011, 2011, 542043. doi:10.4061/2011/542043.

16. Stiller, C.; Viktorsson, K.; Paz Gomero, E.; Haag, P.; Arapi, V.; Kaminskyy, V.O.; Kamali, C.; De Petris, L.; Ekman, S.; Lewensohn, R.; et al.. Detection of Tumor-Associated Membrane Receptors on Extracellular Vesicles from Non-Small Cell Lung Cancer Patients via Immuno-PCR. Cancers 2021, 13. doi:10.3390/cancers13040922.

17. Cabrera-Andrade, A.; López-Cortés, A.; Muñoz, M.J.; Jaramillo-Koupermann, G.; Rodriguez, O.; Leone, P.E.; Paz-y Miño, C. Association of genetic variants of membrane receptors related to recognition and induction of immune response with Helicobacter pylori infection in Ecuadorian individuals. International Journal of Immunogenetics 2014, 41, 281–288. doi:10.1111/iji.12118.

18. Fairbanks, G.; Patel, V.P.; Dino, J.E. Biochemistry of ATP-dependent red cell membrane shape change. Scandinavian Journal of Clinical and Laboratory Investigation. Supplementum 1981, 156, 139–144. doi:10.3109/00365518109097446.

19. Alhanaty, E.; Sheetz, M.P. Cell membrane shape control–effects of chloromethyl ketone peptides. Blood 1984, 63, 1203–1208.

20. Alimohamadi, H.; Smith, A.S.; Nowak, R.B.; Fowler, V.M.; Rangamani, P. Non-uniform distribution of myosin-mediated forces governs red blood cell membrane curvature through tension modulation. PLoS computational biology 2020, 16, e1007890. doi:10.1371/journal.pcbi.1007890.

21. Sakamoto, K.; Morishita, T.; Aburai, K.; Ito, D.; Imura, T.; Sakai, K.; Abe, M.; Nakase, I.; Futaki, S.; Sakai, H. Direct entry of cell-penetrating peptide can be controlled by maneuvering the membrane curvature. Scientific Reports 2021, 11, 31. doi:10.1038/s41598-020-79518-1.

22. Weber, G.; Bianciardi, G.; Toti, P. Circulating platelets plasma-membrane of normocholesterolemic and hypercholesterolemic rabbits tested with aspirin: a freeze etching study of the platelet plasma-membrane “protuberances”. Pharmacological Research Communications 1980, 12, 49–55. doi:10.1016/s0031-6989(80)80062-2.

23. Hennig, T.; O’Hare, P. Viruses and the nuclear envelope. Current opinion in cell biology 2015, 34, 113–121.

24. Chow, K.H.; Factor, R.E.; Ullman, K.S. The nuclear envelope environment and its cancer connections. Nature Reviews Cancer 2012, 12, 196.

25. Bkaily, G.; Avedanian, L.; Al-Khoury, J.; Provost, C.; Nader, M.; D’Orléans-Juste, P.; Jacques, D. Nuclear membrane receptors for ET-1 in cardiovascular function. American Journal of Physiology. Regulatory, Integrative and Comparative Physiology 2011, 300, R251–263. doi:10.1152/ajpregu.00736.2009.

26. Hogeboom, G.H.; Schneider, W.C. On the Nuclear Envelope. Science (New York, N.Y.) 1953, 118, 419. doi:10.1126/science.118.3067.419.

27. Gall, J.G. Observations on the nuclear membrane with the electron microscope. Experimental Cell Research 1954, 7, 197–200. doi:10.1016/0014-4827(54)90054-3.

28. Candia, J.; Maunu, R.; Driscoll, M.; Biancotto, A.; Dagur, P.; McCoy, J.P.; Sen, H.N.; Wei, L.; Maritan, A.; Cao, K.; et al.. From Cellular Characteristics to Disease Diagnosis: Uncovering Phenotypes with Supercells. PLoS Computational Biology 2013, 9, e1003215. doi:10.1371/journal.pcbi.1003215.

29. Candia, J.; Banavar, J.R.; Losert, W. Understanding health and disease with multidimensional single-cell methods. Journal of Physics. Condensed Matter: An Institute of Physics Journal 2014, 26, 073102. doi:10.1088/0953-8984/26/7/073102.

30. Zhao, J. Cell individuality: a basic multicellular phenomenon and its role in the pathogenesis of disease. Medical Hypotheses 1995, 44, 400–402. doi:10.1016/0306-9877(95)90267-8.

31. Zhao, J. A liability theory of disease: the foundation of cell population pathology. Medical Hypotheses 1997, 48, 341–346. doi:10.1016/s0306-9877(97)90104-3.

32. Orrenius, S. Apoptosis: molecular mechanisms and implications for human disease. Journal of Internal Medicine 1995, 237, 529–536. doi:10.1111/j.1365-2796.1995.tb00881.x.

33. Prame Kumar, K.; Nicholls, A.J.; Wong, C.H.Y. Partners in crime: neutrophils and monocytes/macrophages in inflammation and disease. Cell and Tissue Research 2018, 371, 551–565. doi:10.1007/s00441-017-2753-2.

34. Lombard, J. Once upon a time the cell membranes: 175 years of cell boundary research. Biology Direct 2014, 9. doi:10.1186/s13062-014-0032-7.

35. Haralick, R.M.; Shapiro, L.G. Computer and Robot Vision, 1st ed.; Addison-Wesley Longman Publishing Co., Inc., 1992.

36. Pal, N.R.; Pal, S.K. A review on image segmentation techniques. Pattern Recognition 1993, 26, 1277–1294. doi:10.1016/0031-3203(93)90135-J.

37. Taghanaki, S.A.; Abhishek, K.; Cohen, J.P.; Cohen-Adad, J.; Hamarneh, G. Deep semantic segmentation of natural and medical images: A review. Artificial Intelligence Review 2021, 54, 137–178.

38. Romera-Paredes, B.; Torr, P.H.S. Recurrent instance segmentation. European conference on computer vision. Springer, 2016, pp. 312–329.

39. Perez, A.; Seyedhosseini, M.; Deerinck, T.; Bushong, E.; Panda, S.; Tasdizen, T.; Ellisman, M. A workflow for the automatic segmentation of organelles in electron microscopy image stacks. Frontiers in Neuroanatomy 2014, 8, 1–13. doi:10.3389/fnana.2014.00126.

40. Wilke, S.; Antonios, J.; Bushong, E.; Badkoobehi, A.; Malek, E.; Hwang, M. Deconstructing complexity: serial block-face electron microscopic analysis of the hippocampal mossy fiber synapse. Journal of Neuroscience 2013, 33, 507–522. doi:10.1523/JNEUROSCI.1600-12.2013.

41. Bohorquez, D.; Samsa, L.; Roholt, A.; Medicetty, S.; Chandra, R.; Liddle, R. An enteroendocrine cell-enteric glia connection revealed by 3D electron microscopy. PLoS ONE 9:e89881 2014, 9, 1–13. doi:10.1371/journal.pone.0089881.

42. Ørting, S.N.; Doyle, A.; Hilten, A.v.; Hirth, M.; Inel, O.; Madan, C.R.; Mavridis, P.; Spiers, H.; Cheplygina, V. A Survey of Crowdsourcing in Medical Image Analysis. Human Computation 2020, 7, 1–26. doi:10.15346/hc.v7i1.1.

43. Schnoor, J. Citizen science. Environmental Science & Technology 2007, 41, 5923–5923. doi:10.1021/es072599+.

44. Spiers, H.; Songhurst, H.; Nightingale, L.; Folter, J.d.; Hutchings, R.; Peddie, C.J.; Weston, A.; Strange, A.; Hindmarsh, S.; Lintott, C.; et al.. Citizen science, cells and CNNs – deep learning for automatic segmentation of the nuclear envelope in electron microscopy data, trained with volunteer segmentations. bioRxiv 2020, p. 2020.07.28.223024. doi:10.1101/2020.07.28.223024.

45. Cireşan, D.C.; Giusti, A.; Gambardella, L.M.; Schmidhuber, J. Mitosis detection in breast cancer histology images with deep neural networks. International Conference on Medical Image Computing and Computer-assisted Intervention (MICCAI). Springer, 2013, pp. 411–418.

46. Urakubo, H.; Bullmann, T.; Kubota, Y.; Oba, S.; Ishii, S. UNI-EM: An Environment for Deep Neural Network-Based Automated Segmentation of Neuronal Electron Microscopic Images. bioRxiv 2019, p. 607366.

47. Liu, J.; Li, W.; Xiao, C.; Hong, B.; Xie, Q.; Han, H. Automatic Detection and Segmentation of Mitochondria from SEM Images using Deep Neural Network. 2018 40th Annual International Conference of the IEEE Engineering in Medicine and Biology Society (EMBC). IEEE, 2018, pp. 628–631.

48. Dorkenwald, S.; Schubert, P.J.; Killinger, M.F.; Urban, G.; Mikula, S.; Svara, F.; Kornfeld, J. Automated synaptic connectivity inference for volume electron microscopy. Nature methods 2017, 14, 435.

49. Konishi, K.; Mimura, M.; Nonaka, T.; Sase, I.; Nishioka, H.; Suga, M. Practical method of cell segmentation in electron microscope image stack using deep convolutional neural network. Microscopy 2019.

50. Caicedo, J.C.; Roth, J.; Goodman, A.; Becker, T.; Karhohs, K.W.; McQuin, C.; Singh, S.; Carpenter, A.E. Evaulation of Deep Learning Strategies for Nucleus Segmentation in Fluorescence Images. IEEE Reviews in Biomedical Engineering 2018, 2, 147–171.

51. Quan, T.M.; Hildebrand, D.G.C.; Jeong, W. FusionNet: A deep fully residual convolutional neural network for image segmentation in connectomics. CoRR 2016, https://abs/1612.05360, [1612.05360].

52. Dinggang Shen, G.W.; Suk, H.I. Deep Learning in Medical Image Analysis. The Annual Review in Biomedical Engineering 2017, 19, 221–248.

53. Antropova N, Huynh BQ, G.M. A deep feature fusion methodology for breast cancer diagnosis demonstrated on three imaging modality datasets. Med Phys. 2017, 44, 5162–5171.

54. Gulshan, V.; Peng, L.; Coram, M.; Stumpe, M.C.; Wu, D.; Narayanaswamy, A.; Venugopalan, S.; Widner, K.; Madams, T.; Cuadros, J.; et al.. Development and Validation of a Deep Learning Algorithm for Detection of Diabetic Retinopathy in Retinal Fundus Photographs. JAMA 2016, 316, 2402–2410. doi:10.1001/jama.2016.17216.

55. Wang, W.; Wang, Y.; Wu, Y.; Lin, T.; Li, S.; Chen, B. Quantification of full left ventricular metrics via deep regression learning with contour-guidance. IEEE Access 2019, 7, 47918–47928.

56. Bosch, C.; Ackels, T.; Pacureanu, A.; Zhang, Y.; Peddie, C.J.; Berning, M.; Rzepka, N.; Zdora, M.C.; Whiteley, I.; Storm, M.; et al.. Functional and multiscale 3D structural investigation of brain tissue through correlative in vivo physiology, synchrotron micro-tomography and volume electron microscopy. bioRxiv 2021, p. 2021.01.13.426503. doi:10.1101/2021.01.13.426503.

57. Heinrich, L.; Bennett, D.; Ackerman, D.; Park, W.; Bogovic, J.; Eckstein, N.; Petruncio, A.; Clements, J.; Xu, C.S.; Funke, J.; others. Automatic whole cell organelle segmentation in volumetric electron microscopy. bioRxiv 2020.

58. Conrad, R.; Narayan, K. CEM500K, a large-scale heterogeneous unlabeled cellular electron microscopy image dataset for deep learning. Elife 2021, 10. doi:10.7554/elife.65894.

59. Karabağ, C.; Jones, M.L.; Peddie, C.J.; Weston, A.E.; Collinson, L.M.; Reyes-Aldasoro, C.C. Semantic segmentation of HeLa cells: An objective comparison between one traditional algorithm and four deep-learning architectures. Plos One 2020, 15, e0230605.

60. Karabağ, C.; Jones, M.L.; Peddie, C.J.; Weston, A.E.; Collinson, L.M.; Reyes-Aldasoro, C.C. Segmentation and Modelling of the Nuclear Envelope of HeLa Cells Imaged with Serial Block Face Scanning Electron Microscopy. Journal of Imaging 2019, 5(9):75.

61. Deerinck, T.J.; Bushong, E.; Thor, A.; Ellisman, M.H. NCMIR - National Center for Microscopy and Imaging Research. NCMIR methods for 3D EM: A new protocol for preparation of biological specimens for serial block-face SEM Microscopy, 2010.

62. Iudin, A.; Korir, P.K.; Salavert-Torres, J.; Kleywegt, G.J.; Patwardhan, A. EMPIAR: a public archive for raw electron microscopy image data. Nature Methods 2016, 13, 387–388. doi:10.1038/nmeth.3806.

63. Otsu, N. A threshold selection method from gray-level histograms. IEEE Transactions on Systems, Man, and Cybernetics 1979, 9, 62–66.

64. Canny, J. A Computational Approach to Edge Detection. IEEE Transactions on Pattern Analysis and Machine Intelligence 1986, 8, 679–698.

65. Vincent, L.; Soille, P. Watersheds in digital spaces: an efficient algorithm based on immersion simulations. IEEE Transactions on Pattern Analysis and Machine Intelligence 1991, 13, 583–598. doi:10.1109/34.87344.

66. Jaccard, P. Étude comparative de la distribution florale dans une portion des Alpes et des Jura. Bulletin del la Société Vaudoise des Sciences Naturelles 1901, 37, 547–579.

67. Lee, J.K.; Enciso, G.A.; Boassa, D.; Chander, C.N.; Lou, T.H.; Pairawan, S.S.; Guo, M.C.; Wan, F.Y.; Ellisman, M.H.; Sütterlin, C.; others. Replication-dependent size reduction precedes differentiation in Chlamydia trachomatis. Nature communications 2018, 9, 1–9.

